# A Multi-Modal AI/ML-based Framework for Protein Conformation Selection and Prediction in Drug Discovery Applications

**DOI:** 10.64898/2026.02.17.706293

**Authors:** Shivangi Gupta, Vineetha Menon, Jerome Baudry

## Abstract

The development of pharmaceutical drugs is a time-intensive and costly process, with more than 90% of drug candidates failing during preclinical or clinical testing. A major challenge lies in accurately predicting protein-ligand interactions, especially given that traditional computational methods often rely on a single protein conformation, failing to capture biologically relevant structural variability. To address this, we present an AI/ML-based multi-modal framework based on Graph Convolutional Network (GCN) that integrates both global and local protein descriptors to classify binding and non-binding conformations more effectively. Global descriptors capture overarching physico-chemical and structural properties of proteins, while local descriptors—such as pharmacophores—provide site-specific information crucial for modeling ligand interactions. Our GCN based approach demonstrates that integrating local and global structural perspectives significantly improves predictive accuracy and robustness. By enabling more reliable protein conformation classification, this work contributes toward scalable, AI-driven drug discovery—an increasingly critical goal in response to global health challenges.

## Introduction

Drug development is a critical research area for chemical scientists and pharmaceutical companies; however, it continues to face significant challenges, including low success rates, high costs, and lengthy development timelines [1]. Traditional drug discovery approaches frequently fail due to inaccurate target selection, inadequate safety profiles, limited therapeutic efficacy, and difficulties in identifying appropriate patient populations [2]. A major contributor to unsuccessful target selection is off-target binding, in which drug candidates unintentionally interact with proteins other than their intended molecular targets. These off-target interactions can diminish drug selectivity and potency and often lead to adverse side effects or toxicity, ultimately contributing to late-stage clinical failures [3].

In the rapidly evolving landscape of pharmaceutical research, computational methods have become a cornerstone of drug discovery and development. Computer-aided drug design (CADD) offers a cost-effective and time-efficient complement to experimental approaches by enabling the prediction of drug behavior, protein–ligand interactions, and pharmacokinetic properties prior to synthesis and experimental validation. Techniques such as molecular modeling, structure–activity relationship analysis, and virtual screening allow researchers to explore vast chemical spaces and make informed decisions early in the drug development pipeline [4] - [7].

Traditional computational drug discovery, however, often relies on docking ligands to a single, static protein conformation, thereby neglecting biologically relevant receptor flexibility and potentially overlooking effective drug candidates. To overcome this limitation, ensemble-based docking strategies have been developed [8], which incorporate multiple protein conformations generated through molecular dynamics (MD) simulations. This approach better captures conformational variability and enables the identification of hidden or transient binding sites that may not be observable in single-structure docking. In ensemble-based workflows, multiple target protein conformations are screened against pre-existing libraries of active compounds and decoys to identify the strongest and most relevant protein–ligand interactions. Although this approach more effectively captures biologically relevant and high-affinity interactions, it is computationally expensive, particularly when multiple target proteins are considered. Moreover, only a small fraction of the sampled conformations—often just a few hundred out of millions—exhibit statistically significant binding activity that is essential for conformational selection, thereby substantially increasing the computational challenge of identifying these rare but critical states and highlights the need for advanced Big Data analytics, particularly artificial intelligence (AI) to effectively process and interpret large-scale datasets.

The increasing availability of large-scale biochemical and structural data has driven the widespread adoption of artificial intelligence (AI) and machine learning (ML) in modern drug discovery, with the aim of improving efficiency, accuracy, and cost-effectiveness across the development pipeline. AI/ML techniques have shown significant promise in enhancing target specificity, mitigating toxicity risks, and accelerating the progression of drug candidates through early discovery and pre-clinical stages [9, 10]. Ensemble-based docking approaches, which evaluate millions of protein–ligand conformations generated through molecular dynamics simulations, produce extremely large and highly imbalanced datasets. Since only a small fraction of these conformations exhibit meaningful binding activity, efficiently identifying the most relevant candidates remains both computationally intensive and analytically complex. While high-performance computing (HPC) resources are essential to manage the scale of these simulations, intelligent data-driven strategies are equally critical for prioritizing biologically significant conformations. To reduce the substantial computational burden typically associated with ensemble docking, this work introduces performance-oriented AI/ML frameworks that integrate seamlessly with HPC environments and ensemble docking outputs to support protein conformation selection and classification. By directly addressing the challenges of extreme class imbalance and limited experimental validation, these frameworks enable the efficient identification of rare, high-value binding conformations. As a result, they significantly improve the accuracy, robustness, and overall reliability of computational drug discovery pipelines.

In this work, we introduce a multi-modal framework that integrates global and local descriptors to improve the classification of protein conformations. Global descriptors capture overall structural and physicochemical properties—such as mass, hydrophobicity, and radius of gyration, while local descriptors, including pharmacophores, provide site-specific information critical for identifying ligand-binding interactions. These complementary feature sets offer a more comprehensive view, enhancing classification performance. Like global descriptor datasets, pharmacophore data suffer from class imbalance, with only a limited number of conformations showing strong binding activity. This imbalance can lead to biased models and poor generalization. To address this, we apply previously established data-driven augmentation techniques [11]-[13] to balance the dataset and improve model reliability. To capture both spatial and statistical dependencies, we construct graph-based representations for each descriptor type: a proximity matrix for pharmacophores (local descriptors) and a Pearson correlation matrix for global descriptors. These graphs are processed using a Graph Convolutional Network (GCN) trained with a contrastive loss function, which improves feature discrimination by pulling together similar conformations and separating dissimilar ones in the embedding space. The resulting graph-level embeddings are then classified using traditional ML algorithms, and a decision fusion strategy aggregates their outputs for improved performance. This integrated approach combines structural insight with advanced graph learning to provide a scalable, interpretable, and effective solution for protein conformation classification, with potential to accelerate AI-driven drug discovery.

## Materials and methods

### 0.1 Dataset Overview

Four proteins were selected to evaluate the effectiveness of our proposed model: ADORA2A (Adenosine A2A Receptor), ADRB2 (Beta-2 Adrenergic Receptor), OPRD1 (Delta Opioid Receptor), and OPRK1 (Kappa Opioid Receptor). These targets were previously studied and reported in our earlier work [11]– [13]. Each protein includes experimentally validated conformations that either (a) bind to ligands (binding conformations) or (b) do not bind to ligands (non-binding conformations), as previously described in [8].

To improve the classification of binding versus non-binding protein conformations, we employed both global and local structural descriptors, including pharmacophoric features specific to ligand-binding sites. Global descriptors capture overall structural and physicochemical properties of a protein conformation and were described in our prior work [11]– [13]. In contrast, local descriptors were derived from pharmacophore features using the DB-PH4 module in MOE (Molecular Operating Environment). These features were extracted within a 6.5 Å radius of the ligand-binding site using the “unified scheme,” which includes hydrogen bond donors (Don), acceptors (Acc), cations (Cat), anions (Ani), aromatic centers (Aro), and hydrophobic centers (Hyd). The MD-derived conformational dataset containing these local descriptors has also been documented and published in [14].

Table 1 summarizes the number of binding and non-binding protein conformations, the total number of conformations, the number of local and global descriptors, and the class imbalance ratio for each target protein used in this study.

**Table 1.** Dataset description.

| Target protein | # of binding protein conformations | # of non-binding protein conformations | Total # of protein conformations | # of local descriptors | # of global descriptors | Class-imbalance ratio |
| --- | --- | --- | --- | --- | --- | --- |
| **ADORA2A**<br>(adenosine receptor A2A) | 850 | 2,150 | 3,000 | 282 | 50 | 3:1 |
| **ADRB2**<br>(β2-adrenergic receptor) | 154 | 2,411 | 2,565 | 542 | 51 | 16:1 |
| **OPRK1**<br>(κ-type opioid receptor) | 137 | 2,862 | 2,999 | 697 | 50 | 21:1 |
| **OPRD1**<br>(δ-type opioid receptor) | 72 | 2,932 | 3,004 | 329 | 51 | 41:1 |

### 0.2 Pearson Correlation Matrix

Our study’s global features—such as protein mass, volume, radius of gyration, hydrophobic surface area, and mobility—are generic to all protein conformations and lack explicit spatial dependency information for GCN input. To address this, we use the Pearson correlation matrix to capture pairwise linear relationships between features, revealing how changes in one relate to another. It measures the linear relationship between two global descriptors. The correlation coefficient *r*_*ab*_, ranging from -1 to 1, quantifies the strength and direction of this relationship, showing how changes in one feature predict changes in another. It is calculated as follows [15]:

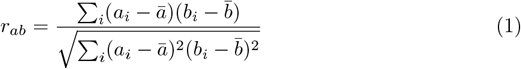

where *a*_*i*_ and *b*_*i*_ are individual feature values and ā, 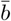 their respective means. A value of +1 indicates a perfect positive linear relationship, -1 a perfect negative linear relationship, and 0 no linear correlation. Values closer to ±1 indicate stronger linear relationships, while those near zero indicate weak or no correlation.

### 0.3 Gaussian Naive Bayes Classifier

The Gaussian Naive Bayes (GB) classifier is a probabilistic supervised learning method based on Bayes’ theorem, assuming feature independence and Gaussian-distributed continuous variables [16]. For each feature, GB estimates the mean (*µ*_*b*_) and variance (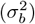) per class *b* to compute the likelihood using:

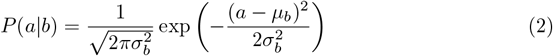

Classification relies on the maximum a posteriori (MAP) estimate derived from Bayes’ theorem:

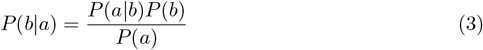

Where *P* (*b*) is the prior, *P* (*a*|*b*) is the likelihood, and *P* (*b*|*a*) is the posterior probability of class *b* given data point *a*. The model predicts the class with the highest posterior probability:

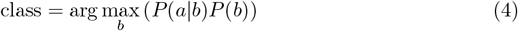

This lightweight and interpretable model is especially effective when the features are conditionally independent and normally distributed [17].

### 0.4 K-Nearest Neighbor

K-Nearest Neighbors (KNN) is a straightforward, non-parametric supervised learning algorithm used for classification and regression. It predicts the label of a new data point *x* by identifying its *K* closest neighbors using the Euclidean distance metric, 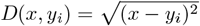, and assigning *x* to the class with the majority among these neighbors. Despite its simplicity, KNN faces limitations such as high memory usage and sensitivity to imbalanced data [18], [19], [20].

### 0.5 Random Forest

Random Forest (RF) is a robust ensemble learning method used for classification and regression. It constructs multiple decision trees during training, and for classification tasks, the final class label is determined by majority voting across these trees, improving accuracy [21].

The training process begins by creating bootstrap samples—random subsets of the training data selected with replacement—ensuring each tree is trained on a unique dataset. At each node, a random subset of features is considered, and the best feature from this subset is selected for splitting. This randomness introduces diversity among trees, reducing overfitting and enhancing generalization [22].

Trees are grown fully without pruning to capture complex data patterns. The best splits are chosen by minimizing the Gini impurity *G*(*M* ), defined as:

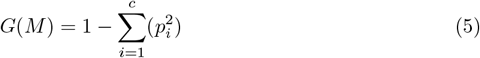

where *G*(*M* ) is the Gini impurity of Node *M, c* is the total number of classes and *p*_*i*_ is the probability of selecting class *i*.

Each tree independently predicts the class of an input sample. For binary classification, the final prediction is the class receiving the majority vote among all trees. For example, if more trees predict Class 1 (binding) than Class 0 (non-binding), the instance is assigned to Class 1. This ensemble approach reduces overfitting, improves accuracy, and increases robustness against noise.

### 0.6 Support Vector Machine

Support Vector Machines (SVMs) are widely used for binary classification by finding the optimal hyperplane that separates two classes in an *n*-dimensional feature space [23]. The hyperplane maximizes the margin, defined as the distance between the hyperplane and the closest points (support vectors) from each class, ensuring reliable class separation. Given a labeled training set {(*x*_1_, *y*_1_), (*x*_2_, *y*_2_), …, (*x*_*n*_, *y*_*n*_)}, where *x*_*n*_ is a feature vector and *y*_*n*_ its class label, the optimal hyperplane satisfies [24]:

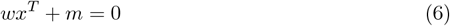

with *w* as the weight vector, *x* the feature vector, and *m* the bias. The classification constraints are:

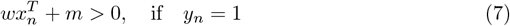

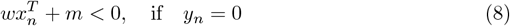

Training adjusts *w* and *m* to maximize the margin 1*/*∥*w*∥^2^, enhancing generalization. For non-linearly separable data, SVM employs kernel functions to implicitly map inputs to higher-dimensional spaces. The linear kernel computes the inner product:

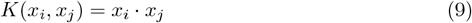

and the decision function is:

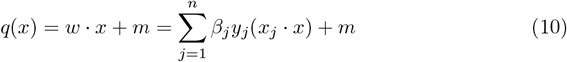

where *β*_*j*_ are Lagrange multipliers [25]. The linear kernel works well when data is linearly separable.

The Radial Basis Function (RBF) kernel extends SVM’s flexibility by using a Gaussian function:

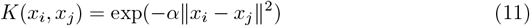

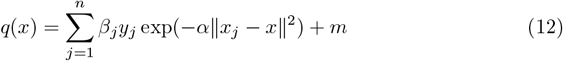

where *α* controls the influence radius of each training point [26]. Low *α* (gamma) yields smoother decision boundaries with risk of underfitting, while high *α* creates tighter boundaries that may overfit noise.

### 0.7 Graph Convolutional Neural Network

Graphs are widely used in domains such as social analysis, bioinformatics, and computer vision to capture structural relationships between data, providing richer insights than analyzing individual data points. A Graph Convolutional Network (GCN) is a deep learning model designed to process graph-structured data, which is inherently non-Euclidean—examples include protein structures, molecules, and social networks. GCNs learn node embeddings by iteratively aggregating information from neighboring nodes, thereby capturing both node features and the overall graph structure [27].

Formally, a graph *G* consists of nodes *N* and edges *E* representing connections between node pairs [28]. In our study, nodes correspond to local or global descriptors, while edges represent spatial distances between local descriptors or linear dependencies expressed as correlation values between global descriptors. The two main components of a GCN are the adjacency matrix *A*, which encodes graph topology, and the node feature matrix *X*, which records node attributes. For our datasets, *X* contains encoded binary vectors for local descriptors and numerical feature values for global descriptors. The GCN aggregates information from neighboring nodes to make predictions. The operation of a GCN layer is described as follows [29]:

- The first step is to add self-loops and normalize the adjacency matrix. Graphs may or may not contain self-loops. Self-loops are included to ensure that each node takes into account its own feature. Equation (13) shows the creation of matrix Ā by adding self-loops to the adjacency matrix *A*, where *E* represents the identity matrix used to incorporate self-connections.

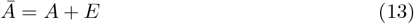

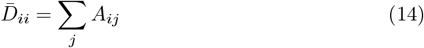

Here, 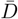 denotes the degree matrix, which can be computed as shown in Equation (14). The normalized adjacency matrix *U* is then calculated using Equation (15).

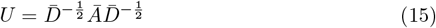

- The next step is node feature aggregation, where each node combines its own feature vector with those of its neighbors to obtain local context. This aggregation process leverages the normalized adjacency matrix *U* and is expressed in Equation (16):

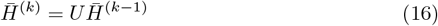

Here, 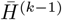 represents node features from the previous layer. This operation successfully propagates and blends information across neighboring nodes, guaranteeing that each node representation is enhanced by its local graph structure.

- The following step is feature transformation, which involves the GCN applying a learnable linear transformation on the aggregated node features. This technique is similar to using a Dense (or Linear) layer in a classical neural network. It allows the model to modify, combine, and enhance the aggregated data to better fit the task at hand. Equation (17) illustrates the computation of the transformed feature matrix:

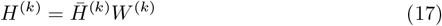

Here, *W* ^(*k*)^ is the learnable weight matrix that determines how the aggregated features are transformed.

- Lastly, a non-linearity, such as *ReLU*, is applied element-by-element to the transformed features, as indicated in Equation (18). This activation function introduces nonlinearity to the model, allowing it to capture complicated patterns and relationships in the same way that hidden layers in traditional neural networks do.

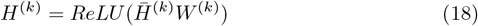

The GCN model in this study consists of two graph convolutional layers followed by a mean pooling layer, which aggregates node features by averaging them to produce a fixed-size graph-level embedding that captures the overall structure and feature distribution.

To enhance the embeddings’ discriminative power, the GCN is trained using a contrastive loss function that pulls embeddings of similar graphs closer while pushing dissimilar ones apart. The loss is defined as follows:

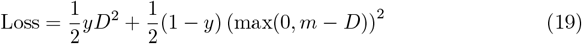

where *D* = ∥*e*_*i*_ − *e*_*j*_∥^2^ is the Euclidean distance between embeddings *e*_*i*_ and *e*_*j*_, and *y* indicates pair similarity (*y* = 1 for similar pairs—both binding or non-binding—and *y* = 0 for dissimilar pairs). The margin *m* sets the minimum distance for negative pairs. The first term pulls positive pairs together, while the second pushes negative pairs apart if closer than *m*.

### 0.8 Decision Fusion

In this study, we provide a decision fusion strategy that improves the detection of binding and non-binding protein conformations while also enhancing the AI/ML model’s capacity to distinguish between them. We propose a method that leverages the predictive power of four machine learning techniques which operate on embeddings produced by a unique dual Graph Convolution Network (GCN) model. This dual GCN model consists of two separately trained GCN on a dataset that contains local spatial and global linear relationship respectively, allowing it to capture a more complete description of the data.

Let us define the prediction results obtained from the four machine learning models—Gaussian Naïve Bayes (GB), K-Nearest Neighbor (KNN), Random Forest (RF), and Support Vector Machine (SVM)—applied to GCN embeddings of local and global descriptors as follows: *L*_*GB*_, *L*_*KNN*_, *L*_*RF*_ and *L*_*SV M*_ represent the predictions obtained using embeddings of GCN model trained on local descriptors and *G*_*GB*_, *G*_*KNN*_, *G*_*RF*_ and *G*_*SV M*_ represent the predictions obtained using embeddings of GCN model trained on global descriptors. After obtaining the individual predictions from each dataset, the cumulative prediction score is computed by summing all the predictions, as defined by the following equation (19).

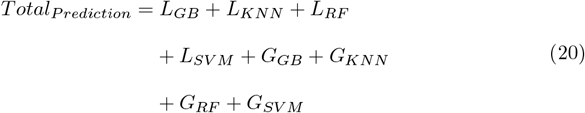

where *Total*_*P rediction*_ is used to compute the final classification results. Conformations with a cumulative score of at least 3 are allocated to Class ‘1’, indicating binding protein conformation, while those with a cumulative score of no more than 5 are assigned to Class ‘0’, indicating non-binding protein conformation.

### 0.9 Evaluation metrics

The confusion matrix and its derived metrics—such as accuracy, sensitivity, and specificity—are widely used to evaluate the performance of machine learning classifiers. For binary classification of binding vs. non-binding protein conformations, the confusion matrix includes four outcomes [11]:

- True Positive (TP): Correctly predicted binding conformations (Class 1).
- False Positive (FP): Non-binding conformations incorrectly predicted as binding.
- False Negative (FN): Binding conformations incorrectly predicted as non-binding.
- True Negative (TN): Correctly predicted non-binding conformations (Class 0).

The accuracy measures the overall proportion of correct predictions:

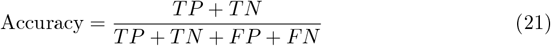

Sensitivity quantifies the model’s ability to correctly identify binding conformations:

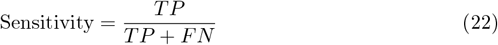

Specificity quantifies the model’s ability to correctly identify non-binding conformations:

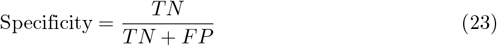

## 0.10 Enrichment Ratio

To validate the proposed AI/ML framework, true positive (TP) and false negative (FN) predictions were used to compute the enrichment ratio. As outlined in our previous work [11], a baseline enrichment ratio—representing the expected performance under random selection—was calculated for comparison. This metric provides a benchmark to assess how effectively the AI/ML framework improves prediction performance relative to random selection of protein conformations.

Table 2 summarizes the maximum Base Enrichment ratios computed for each target protein.

**Table 2.** Maximum Base Enrichment ratios calculated for each target proteins.

| Protein | Maximum Enrichment Ratio |
| --- | --- |
| ADORA2A | $\sim 9$ |
| ADRB2 | $\sim 5.4$ |
| OPRD1 | $\sim 2$ |
| OPRK1 | $\sim 5.5$ |

### 0.11 Proposed Work

This study introduces a dual-GCN framework that integrates both local and global features for protein conformation prediction. One GCN is trained on pharmacophore-based local descriptors to capture spatial interactions at binding sites, while the other learns from global descriptors to model broader structural patterns. Each GCN is optimized using a contrastive loss to enhance the separation between binding and non-binding conformations in the embedding space. The resulting embeddings are then fed into four traditional machine learning classifiers, whose outputs are combined through decision fusion to yield the final prediction. An enrichment ratio framework is applied to validate binding conformations in test proteins. This approach leverages the representational strength of GCNs and the predictive power of classical models to improve protein conformation selection. An overview of the method is shown in Fig 1.

**Fig 1.**
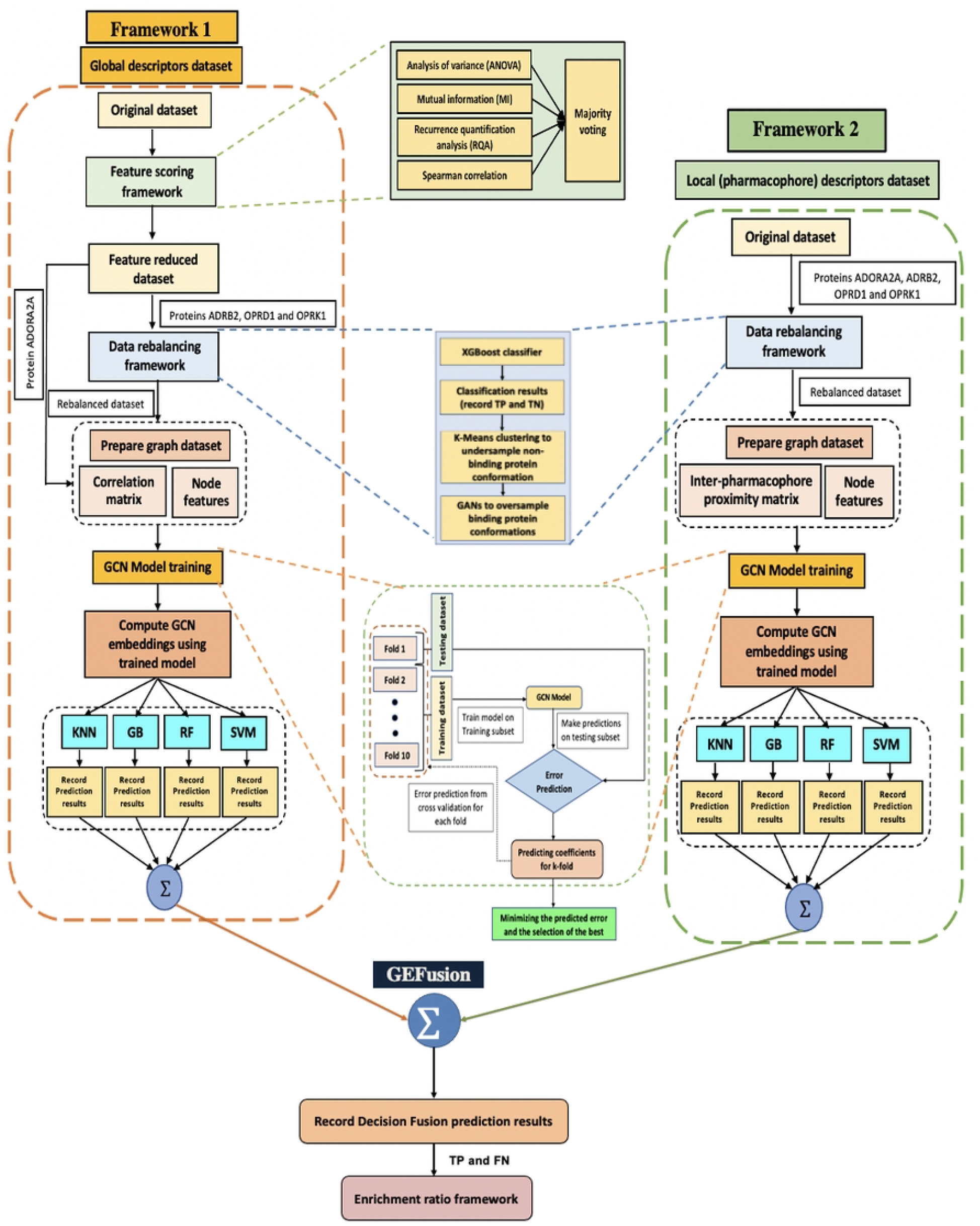
Bold the figure title. Figure caption text here, please use this space for the figure panel descriptions instead of using subfigure commands. A: Lorem ipsum dolor sit amet. B: Consectetur adipiscing elit.

- The summary of Framework 1, which handles the global descriptor data, is as follows:
  - It begins with the original dataset, which is then refined using the feature scoring method described earlier.
  - The ML-based feature selection and scoring framework from our prior work [12], [13], is used to identify the most significant protein characteristics. Specifically, four methods are used: Analysis of Variance (ANOVA) [30], Mutual Information (MI) [31], Recurrence Quantification Analysis (RQA) [32], and Spearman Correlation [33]. Each method independently ranks the features based on its criteria. These rankings are then integrated using a majority voting strategy, where features that receive the highest possible vote count of 4—indicating consensus across all methods—are selected. The resulting subset of features is used to construct a new, optimized dataset for further analysis. This framework, along with the selected features for each target protein—ADORA2A, ADRB2, OPRD1, and OPRK1—has been previously described and published in [12].
  - To address the high class imbalance observed in the proteins ADRB2, OPRK1, and OPRD1, a multi-step data re-balancing approach was implemented as detailed in [12]. First, the Extreme Gradient Boosting (XGBoost) [34] classifier was applied to the feature-reduced dataset to distinguish between non-binding (TN) and binding (TP) protein conformations. Next, K-Means [35]-[37] clustering was performed on the non-binding conformations to retain representative samples while eliminating redundant data, thereby reducing dataset bias and size. Finally, Generative Adversarial Networks (GANs) were employed to augment the minority class by generating additional binding conformation samples, achieving balanced class representation without increasing the overall dataset size. For the ADORA2A protein, this re-balancing step was omitted due to its low class imbalance, which did not necessitate adjustment for the GCN model.
  - In parallel with the re-balancing procedure, a Pearson correlation matrix was constructed to capture pairwise linear relationships among the global features. This analysis provides insight into how changes in one feature relate to changes in another, enhancing understanding of feature interactions. Following this, a graph dataset was prepared for input into the GCN model. Here, global features correspond to nodes, the Pearson correlation coefficients serve as edge weights, and the numerical values of the features are assigned as node attributes.
  - The GCN model was then trained on this graph dataset using 10-fold cross-validation to ensure reliable and generalizable performance. Early stopping was incorporated to prevent overfitting by monitoring the validation loss and halting training when no further improvement was observed. The model with the best validation results was saved and used to generate the final graph-level embeddings.
  - These graph-level embeddings, derived from the optimized GCN model, were subsequently fed into four traditional machine learning classifiers — Gaussian Naïve Bayes (GB), K-Nearest Neighbor (KNN), Random Forest (RF), and Support Vector Machine (SVM) — to assess and compare their classification performances.
- The summary of Framework 2, which processes local descriptor (pharmacophore) data, is as follows:
  - The framework begins by applying the multi-step re-balancing technique described in [12]. Unlike the global descriptor dataset, the original local descriptor data is excluded from the feature scoring process due to its high sparsity. Sparse data poses challenges for GCN training because limited and uneven feature availability hampers the aggregation of meaningful patterns across nodes. This issue is exacerbated by conformations containing only one or two pharmacophores in close proximity. Removing such features based on relevance risks losing critical local information necessary for capturing conformation-specific interactions. Therefore, to preserve essential local information, the original local descriptor dataset bypasses the feature scoring framework.
  - To address class imbalance, the previously described multi-step data re-balancing method is applied. Initially, the feature-reduced dataset is classified with XGBoost to distinguish binding (TP) from non-binding (TN) conformations. K-Means clustering is then used on the TN samples to retain representative instances and reduce redundancy. Finally, GANs augment the minority class by generating additional binding conformations, balancing the classes without increasing overall dataset size.
  - Next, a graph dataset is prepared for input to the GCN model, where local features serve as nodes, and the distances between them define edge weights. Each node is further characterized by attributes including an encoded binary vector, radius, and frequency corresponding to the respective local feature.
  - The graph dataset is processed using the GCN model with 10-fold cross-validation and early stopping to ensure stable, reliable performance and prevent overfitting. The model exhibiting the best validation performance is selected to generate the final graph-level embeddings, which are then input into four machine learning classifiers for evaluation and classification performance recording.
  - GEFusion (Graph Embedding Fusion), a decision-level fusion strategy, is applied to combine the classification results from Framework 1 and Framework 2. This integration enables the unique identification of the total number of probable binding and non-binding protein conformations.
- Finally, enrichment ratios are calculated using True Positives (TP)—correctly predicted binding conformations—and False Negatives (FN)—incorrectly predicted non-binding conformations—based on the outcomes of the GEFusion decision strategy.

## Results and Discussion

### 0.12 Computational Evaluation on ADORA2A Dataset

Table 3 presents the classification performance of the ADORA2A protein across varying training sizes, evaluated using the proposed GEFusion methodology. Notably, the model achieves its highest performance—measured by accuracy, sensitivity, and specificity—when trained on 40% of the dataset. The results for this training size, including the performance of individual classifiers applied to both local and global embeddings, are summarized in Table 4

**Table 3.** Classification performance (in %) across different training sizes for protein ADORA2A.

| Training Size | Accuracy (%) | Sensitivity (%) | Specificity (%) |
| --- | --- | --- | --- |
| 0.1 | 90.93 | 88.09 | 92.05 |
| 0.2 | 91.88 | 84.66 | 94.72 |
| 0.3 | 88.52 | 72.90 | 94.53 |
| 0.4 | 94.94 | 93.24 | 95.61 |
| 0.5 | 89.00 | 70.05 | 96.22 |
| 0.6 | 87.83 | 68.96 | 95.14 |
| 0.7 | 92.00 | 85.48 | 94.48 |
| 0.8 | 89.33 | 68.45 | 97.45 |

**Table 4.**
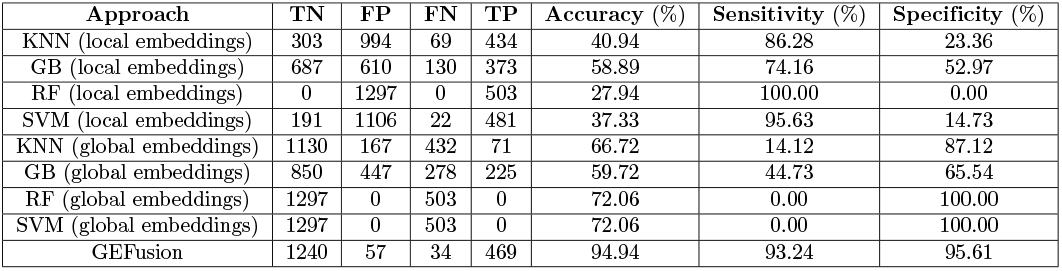
Classification performance for 40% training size for protein ADORA2A.

| Approach | TN | FP | FN | TP | Accuracy (%) | Sensitivity (%) | Specificity (%) |
| --- | --- | --- | --- | --- | --- | --- | --- |
| KNN (local embeddings) | 303 | 994 | 69 | 434 | 40.94 | 86.28 | 23.36 |
| GB (local embeddings) | 687 | 610 | 130 | 373 | 58.89 | 74.16 | 52.97 |
| RF (local embeddings) | 0 | 1297 | 0 | 503 | 27.94 | 100.00 | 0.00 |
| SVM (local embeddings) | 191 | 1106 | 22 | 481 | 37.33 | 95.63 | 14.73 |
| KNN (global embeddings) | 1130 | 167 | 432 | 71 | 66.72 | 14.12 | 87.12 |
| GB (global embeddings) | 850 | 447 | 278 | 225 | 59.72 | 44.73 | 65.54 |
| RF (global embeddings) | 1297 | 0 | 503 | 0 | 72.06 | 0.00 | 100.00 |
| SVM (global embeddings) | 1297 | 0 | 503 | 0 | 72.06 | 0.00 | 100.00 |
| GEFusion | 1240 | 57 | 34 | 469 | 94.94 | 93.24 | 95.61 |

From Table 4, it is evident that while the overall framework delivers strong classification performance, some individual models exhibited extreme predictive behavior. For instance, the Random Forest (RF) model trained on local embeddings classified all conformations as binding, whereas both RF and Support Vector Machine (SVM) models trained on global embeddings predicted all conformations as non-binding. These individual biases are effectively mitigated by the framework’s voting-based ensemble mechanism, which combines predictions from multiple models. This ensemble strategy reduces the impact of the outlier behavior of any single model, resulting in more balanced and reliable classification results.

As shown in Table 5, our proposed methodology demonstrated strong performance not only in overall model accuracy but also in achieving a high enrichment ratio.

**Table 5.** Enrichment ratios of protein ADORA2A on a training size of 40%.

| Approach | Maxima | Filter | % of Data | Minima | Filter | % of Data |
| --- | --- | --- | --- | --- | --- | --- |
| GEFusion | 13.67 | Filter A | 0.5% | 13.07 | Filter A | 1.0% |

### 0.13 Computational Evaluation on ADRB2 Dataset

Table 6 presents the classification performance of the proposed approach across various training sizes for the ADRB2 protein. Among these, the 40% training size yields the best overall performance, achieving higher accuracy, sensitivity, and specificity. Detailed results for this optimal training size—including the performance of the proposed framework and individual classifiers utilizing local and global embeddings—are provided in Table 7.

**Table 6.** Classification performance (in %) across different training sizes for protein ADRB2.

| Training Size | Accuracy (%) | Sensitivity (%) | Specificity (%) |
| --- | --- | --- | --- |
| 0.1 | 84.11 | 64.08 | 85.42 |
| 0.2 | 88.35 | 65.63 | 89.86 |
| 0.3 | 88.47 | 64.29 | 90.08 |
| 0.4 | 92.14 | 80.61 | 92.92 |
| 0.5 | 93.06 | 42.68 | 96.50 |
| 0.6 | 90.55 | 56.06 | 92.92 |
| 0.7 | 86.49 | 66.00 | 87.92 |
| 0.8 | 93.37 | 8.82 | 99.37 |

**Table 7.** Classification performance for 40% training size for protein ADRB2.

| Approach | TN | FP | FN | TP | Accuracy (%) | Sensitivity (%) | Specificity (%) |
| --- | --- | --- | --- | --- | --- | --- | --- |
| KNN (local embeddings) | 634 | 807 | 49 | 49 | 44.38 | 50.00 | 44.00 |
| GB (local embeddings) | 355 | 1086 | 22 | 76 | 28.01 | 77.55 | 24.64 |
| RF (local embeddings) | 348 | 1093 | 22 | 76 | 27.55 | 77.55 | 24.15 |
| SVM (local embeddings) | 260 | 1181 | 17 | 81 | 22.16 | 82.65 | 18.04 |
| KNN (global embeddings) | 978 | 463 | 60 | 38 | 66.02 | 38.78 | 67.87 |
| GB (global embeddings) | 1000 | 441 | 62 | 36 | 67.32 | 36.73 | 69.40 |
| RF (global embeddings) | 1376 | 65 | 87 | 11 | 92.06 | 11.22 | 95.49 |
| SVM (global embeddings) | 1372 | 69 | 92 | 6 | 89.54 | 6.12 | 95.21 |
| GEFusion | 1339 | 102 | 19 | 79 | 92.14 | 80.61 | 92.92 |

Table 8 provides an overview of the derived enrichment ratios based on the decision model’s prediction outcomes (TP and FN), serving as a validation measure for the predictions generated by our proposed framework.

**Table 8.** Enrichment ratios of protein ADRB2 on a training size of 40%.

| Approach | Maxima | Filter | % of Data | Minima | Filter | % of Data |
| --- | --- | --- | --- | --- | --- | --- |
| GEFusion | 33.71 | Filter A | 0.5% | 28.10 | Filter C | 1.0% |

### 0.14 Computational Evaluation on OPRD1 Dataset

Table 9 presents the classification performance of the protein OPRD1 across various training sizes using our AI/ML-based multi-modal framework for protein conformation selection and prediction. The best overall performance—measured by accuracy, sensitivity, and specificity—is achieved with a training size of 10%. Table 10 further details the classification results of the proposed framework and the individual performance of classifiers utilizing local and global embeddings at this optimal training size.

**Table 9.**
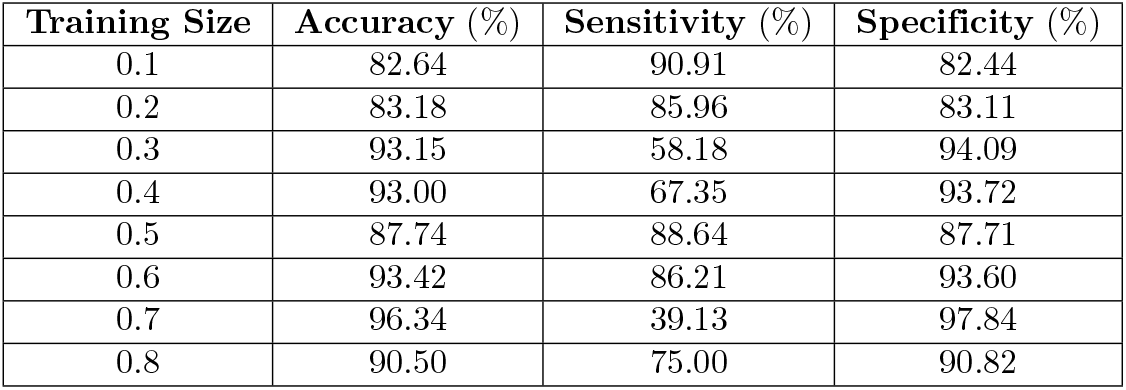
Classification performance (in %) across different training sizes for protein OPRD1.

| Training Size | Accuracy (%) | Sensitivity (%) | Specificity (%) |
| --- | --- | --- | --- |
| 0.1 | 82.64 | 90.91 | 82.44 |
| 0.2 | 83.18 | 85.96 | 83.11 |
| 0.3 | 93.15 | 58.18 | 94.09 |
| 0.4 | 93.00 | 67.35 | 93.72 |
| 0.5 | 87.74 | 88.64 | 87.71 |
| 0.6 | 93.42 | 86.21 | 93.60 |
| 0.7 | 96.34 | 39.13 | 97.84 |
| 0.8 | 90.50 | 75.00 | 90.82 |

**Table 10.** Classification performance for 10% training size for protein OPRD1.

| Approach | TN | FP | FN | TP | Accuracy (%) | Sensitivity (%) | Specificity (%) |
| --- | --- | --- | --- | --- | --- | --- | --- |
| KNN (local embeddings) | 1749 | 887 | 20 | 46 | 66.43 | 69.70 | 66.35 |
| GB (local embeddings) | 1783 | 853 | 26 | 40 | 67.47 | 60.61 | 67.64 |
| RF (local embeddings) | 1598 | 1038 | 12 | 54 | 61.14 | 81.82 | 60.62 |
| SVM (local embeddings) | 2087 | 549 | 33 | 33 | 78.46 | 50.00 | 79.17 |
| KNN (global embeddings) | 1308 | 1328 | 18 | 48 | 50.19 | 72.73 | 49.62 |
| GB (global embeddings) | 1225 | 1411 | 18 | 38 | 47.30 | 67.86 | 46.47 |
| RF (global embeddings) | 1558 | 1078 | 23 | 43 | 59.25 | 65.15 | 59.10 |
| SVM (global embeddings) | 1025 | 1611 | 11 | 55 | 39.97 | 83.33 | 38.88 |
| GEFusion | 2173 | 463 | 6 | 60 | 82.64 | 90.91 | 82.44 |

Table 11 presents the resulting enrichment ratios derived from the predictions made by the GEFusion decision strategy, further supporting the validity and effectiveness of the proposed framework.

**Table 11.** Enrichment ratios of protein OPRD1 on a training size of 10%.

| Approach | Maxima | Filter | % of Data | Minima | Filter | % of Data |
| --- | --- | --- | --- | --- | --- | --- |
| GEFusion | 38.33 | Filter A | 1.0% | 32.38 | Filter B | 0.5% |

### 0.15 Computational Evaluation on OPRK1 Dataset

Table 12 summarizes the classification performance across various training sizes for the protein OPRK1 using the proposed AI/ML-based multimodal framework for protein conformation selection and prediction. The highest overall performance—reflected by improved accuracy, sensitivity, and specificity—was achieved with a training size of 30%. Table 13 presents the classification results at this training size, including the performance of individual classifiers applied to both local and global embeddings.

**Table 12.** Classification performance (in %) across different training sizes for protein OPRK1.

| Training Size | Accuracy (%) | Sensitivity (%) | Specificity (%) |
| --- | --- | --- | --- |
| 0.1 | 83.33 | 64.00 | 84.27 |
| 0.2 | 86.04 | 62.07 | 87.25 |
| 0.3 | 73.08 | 81.19 | 72.67 |
| 0.4 | 77.88 | 79.31 | 77.80 |
| 0.5 | 84.66 | 70.27 | 85.40 |
| 0.6 | 92.17 | 53.33 | 94.21 |
| 0.7 | 78.67 | 73.47 | 78.97 |
| 0.8 | 84.17 | 72.73 | 84.60 |

**Table 13.** Classification performance for 30% training size for protein OPRK1.

| Approach | TN | FP | FN | TP | Accuracy (%) | Sensitivity (%) | Specificity (%) |
| --- | --- | --- | --- | --- | --- | --- | --- |
| KNN (local embeddings) | 1005 | 993 | 52 | 49 | 50.21 | 48.51 | 50.30 |
| GB (local embeddings) | 1295 | 703 | 61 | 40 | 63.60 | 39.60 | 64.81 |
| RF (local embeddings) | 0 | 1998 | 0 | 101 | 4.81 | 100.00 | 0.00 |
| SVM (local embeddings) | 745 | 1253 | 29 | 72 | 38.92 | 71.29 | 37.29 |
| KNN (global embeddings) | 1066 | 932 | 46 | 55 | 53.41 | 54.46 | 53.35 |
| GB (global embeddings) | 1317 | 681 | 52 | 58 | 60.58 | 48.51 | 65.92 |
| RF (global embeddings) | 1273 | 725 | 54 | 47 | 62.89 | 46.53 | 63.71 |
| SVM (global embeddings) | 987 | 1011 | 38 | 63 | 50.02 | 62.38 | 49.40 |
| GEFusion | 1452 | 546 | 19 | 82 | 73.08 | 81.19 | 72.67 |

As shown in Table 14, our proposed methodology demonstrated strong performance not only in overall model accuracy but also in achieving a high enrichment ratio.

**Table 14.**
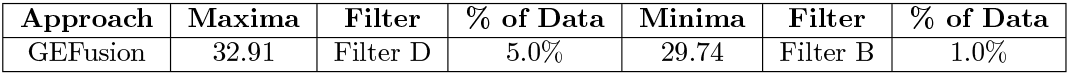
Enrichment ratios of protein OPRK1 on a training size of 30%.

| Approach | Maxima | Filter | % of Data | Minima | Filter | % of Data |
| --- | --- | --- | --- | --- | --- | --- |
| GEFusion | 32.91 | Filter D | 5.0% | 29.74 | Filter B | 1.0% |

From Tables 3, 6, 9, and 12, it is observed that the model consistently performs well across all training sizes for protein ADORA2A. However, for proteins ADRB2, OPRK1, and OPRD1, better overall performance is achieved with smaller training sizes. This trend likely stems from higher class imbalance ratios in these datasets. At larger training sizes, the increased proportion of synthetic binding conformation data, coupled with inherent data sparsity, challenges the GCN model’s ability to distinguish binding from non-binding classes, thus affecting the framework’s predictive accuracy.

The enrichment ratios shown in Tables 5, 8, 11, and 14 demonstrate that binding conformations can be identified with significantly higher accuracy compared to random sampling, as evidenced by the baseline results in Table 2. In particular, the highest enrichment is achieved when selecting the top 0.5% to 1% of the binding conformations, indicating that the most predictive and biologically relevant data points are concentrated among the candidates ranked highest.

## Conclusion

This work presented an AI/ML-based multi-modal framework designed to improve the detection of both binding and non-binding protein conformations by integrating local and global structural information. The proposed approach leverages Graph Convolutional Networks (GCNs) trained with a contrastive loss function to capture meaningful connectivity patterns within protein structures. The GCN learns discriminative graph embeddings that effectively group binding conformations while separating non-binding ones. These embeddings are then processed by traditional machine learning classifiers, and their outputs are combined using a decision fusion strategy to enhance overall classification performance.

The study demonstrated that incorporating both global descriptors—reflecting protein-level structural and stability features—and local descriptors—such as pharmacophores that encode site-specific binding information—provides complementary insights that significantly improve conformation classification. Additionally, the use of graph embeddings learned via contrastive learning offers a more compact and noise-resistant data representation than conventional graph classification methods. These embeddings preserve essential structural relationships while facilitating a clearer distinction between classes.

Overall, the integration of multi-modal data, GCN-based embedding learning, and ensemble decision-making contributes to a robust and generalizable framework capable of accurately identifying protein binding states. This methodology holds promise for advancing structure-based protein function prediction and accelerating drug discovery workflows.

## Acknowledgments

The authors gratefully acknowledge Dr. Armin Ahmadi for providing the Pharmacophore dataset used in this study.

